# Spherical Echo-Planar Time-resolved Imaging (sEPTI) for rapid 3D quantitative T2* and Susceptibility imaging

**DOI:** 10.1101/2024.02.21.581459

**Authors:** Nan Wang, Congyu Liao, Xiaozhi Cao, Mark Nishimura, Yannick WE Brackenier, Mahmut Yurt, Mengze Gao, Daniel Abraham, Cagan Alkan, Siddharth Srinivasan Iyer, Zihan Zhou, Adam Kerr, Justin P. Haldar, Kawin Setsompop

**Author notes:** **Correspondence** Kawin Setsompop, PhD Department of Radiology Stanford University Stanford, CA, United States (94305).

## Abstract

**Purpose:** To develop a 3D spherical EPTI (sEPTI) acquisition and a comprehensive reconstruction pipeline for rapid high-quality whole-brain submillimeter T2* and QSM quantification.

**Methods:** For the sEPTI acquisition, spherical k-space coverage is utilized with variable echo-spacing and maximum k_x_ ramp-sampling to improve efficiency and incoherency when compared to existing EPTI approaches. For reconstruction, an iterative rank-shrinking B_0_ estimation and odd-even high-order phase correction algorithms were incorporated into the reconstruction to better mitigate artifacts from field imperfections. A physics-informed unrolled network was utilized to boost the SNR, where 1-mm and 0.75-mm isotropic whole-brain imaging were performed in 45 and 90 seconds, respectively. These protocols were validated through simulations, phantom, and in vivo experiments. Ten healthy subjects were recruited to provide sufficient data for the unrolled network. The entire pipeline was validated on additional 5 healthy subjects where different EPTI sampling approaches were compared. Two additional pediatric patients with epilepsy were recruited to demonstrate the generalizability of the unrolled reconstruction.

**Results:** sEPTI achieved 1.4 × faster imaging with improved image quality and quantitative map precision compared to existing EPTI approaches. The B0 update and the phase correction provide improved reconstruction performance with lower artifacts. The unrolled network boosted the SNR, achieving high-quality T2* and QSM quantification with single average data. High-quality reconstruction was also obtained in the pediatric patient using this network.

**Conclusion:** sEPTI achieved whole-brain distortion-free multi-echo imaging and T2* and QSM quantification at 0.75 mm in 90 seconds which has the potential to be useful for wide clinical applications.

## 1 Introduction

Quantitative MRI has been used to evaluate a large number of structural and functional characteristics in neuroimaging, with promising clinical and neuroscientific applications^1–3^. For example, T2* mapping and quantitative susceptibility mapping (QSM) have played crucial roles for iron quantification in the brain^4,5^. Iron is critical in numerous neurological activities, including oxygen transport, neurotransmitter synthesis, and myelination^6,7^; abnormal iron accumulation is also associated with various neurological diseases^8–11^. Both R2* (the reciprocal of T2*) and QSM values are proportionately correlated with iron concentration in vivo^4,12^. Additionally, combining T2* and QSM can provide complementary information on myelin content^13,14^. T2* and QSM have been applied to the studies of brain development^15^, functional activity^16^, iron accumulation during aging^17^, and the characterization of neurodegenerative diseases^18^, with promising outcomes.

Multi-echo gradient echo (GRE) sequence is the standard approach for T2* and QSM quantification^19^, but suffers from long scan time. T2* and QSM require acquisition at long echo times (TE up to 30-50 ms at 3T) to generate sufficient magnitude or phase contrast. Particularly for QSM, data acquisition with full 3D brain coverage at a voxel size ≤ 1 mm^3^ is desired to avoid severe partial volume effects^20^. To fulfill these requirements, a typical clinical multi-echo GRE protocol takes 5 to 10 minutes, which is relatively long and inapplicable to some populations. Also, to alleviate the artifacts from odd-even phase differences from switching gradient with bipolarity acquisition ^21^, multi-echo GRE usually uses monopolar acquisition with a flyback gradient between echoes^20^, which reduces the acquisition efficiency and increases the echo spacing.

Echo planar imaging (EPI) acquisition can be a substitute for multi-echo GRE and can improve the acquisition speed for T2* mapping^22^ and QSM^23^, to just tens of seconds. However, EPI suffers from field-inhomogeneity-induced geometric distortion and T2/T2* blurring-induced resolution loss. Moreover, EPI is built on dual polarity sampling with fast gradient switching, which is inevitably subject to artifacts related to odd-even phase differences^21^. These artifacts can be particularly significant when using a higher slew rate and/or a large amount of ramp sampling for improved efficiency.

Advanced techniques have been developed for faster T2* and QSM quantification^24–32^. For example, STrategically Acquired Gradient Echo (STAGE)^24^ was developed to provide T1, T2*, and QSM mapping with upper brain coverage (128 cm) and 0.67 × 1.33 × 2.0 mm^3^ resolution within 5 minutes. Blip up-down acquisition for spin- and gradient-echo imaging (BUDA-SAGE)^25^ used rapid EPI sampling to achieve T2, T2*, and QSM separation with upper brain coverage of 1 × 1 × 2 mm^3^ within 45 seconds. Here, EPI distortion is mitigated using blip-up-down sampling, with T2* blurring being reduced via higher parallel imaging acceleration per EPI-shot. Lastly, Echo planar time-resolved imaging (EPTI)^26–31^ was recently developed to provide fast distortion-free and blurring-free multi-echo acquisition, where T2* maps at 1 mm isotropic resolution with upper brain coverage can be obtained within 1 minute^31^. Nonetheless, further improvements in encoding efficiency and speed are still desirable to achieve faster imaging, with larger coverage and higher resolution. Moreover, residual odd-even field artifacts will limit the usage of higher slew rate and/or more ramp sampling for improved acquisition efficiency. SNR is another challenge for T2* and QSM due to the rapidly decreasing signal intensity at long TEs. Data averaging is usually needed for sufficient SNR, resulting in a doubling or tripling of the scan time.

To address the abovementioned limitations, we developed a novel spherical EPTI (sEPTI) technique with a comprehensive reconstruction pipeline for ultrafast and robust quantitative neuroimaging. The innovations include: (1) to reduce the scan time as well as increase the temporal sampling incoherence across k-space, a sEPTI sampling was designed, which traverses a tight spherical k-space coverage with varying echo spacing at different k_yz_-space locations; (2) to obtain high-quality image reconstruction, an iterative rank-shrinking B_0_ (field inhomogeneity) estimation approach was developed to achieve more accurate B_0_ for EPTI reconstruction; (3) a higher-order odd-even phase correction model was implemented and evaluated to mitigate related artifacts; (4) a physics-informed unrolled network^33^ for subspace reconstruction was also incorporated to boost the SNR for single-average acquisition at submillimeter resolution. Putting together, the proposed sEPTI technique achieved whole-head, high-SNR, distortion- and blurring-free quantification of T2* and QSM at 1-mm isotropic resolution in 45 seconds, and at 0.75-mm isotropic resolution in 90 seconds.

## 2 Theory

### 2.1 sEPTI sampling design

#### 2.1.1 Previous EPTI trajectory

EPTI is a Cartesian spatiotemporal encoding technique along TE dimension for distortion-free and blurring-free neuroimaging. A typical k_y_-t sampling pattern for 2D EPTI^26^ is illustrated in Figure 1A. Each EPTI shot covers a certain k_y_ area with a spatiotemporal CAIPI trajectory. After spatiotemporal reconstruction, each echo time produces one distortion-free and blurring-free T2-weighted (with spin echo sequence) or T2*-weighted (with gradient echo sequence) image, allowing further quantitative mapping. This 2D sampling can be directly extended to 3D with full partition sampling along the k_z_ direction, referred to as partition-EPTI. The acquisition time is the product of the number of shots in a partition, the number of partitions, and the repetition time (TR) of each shot.

**Figure 1:**
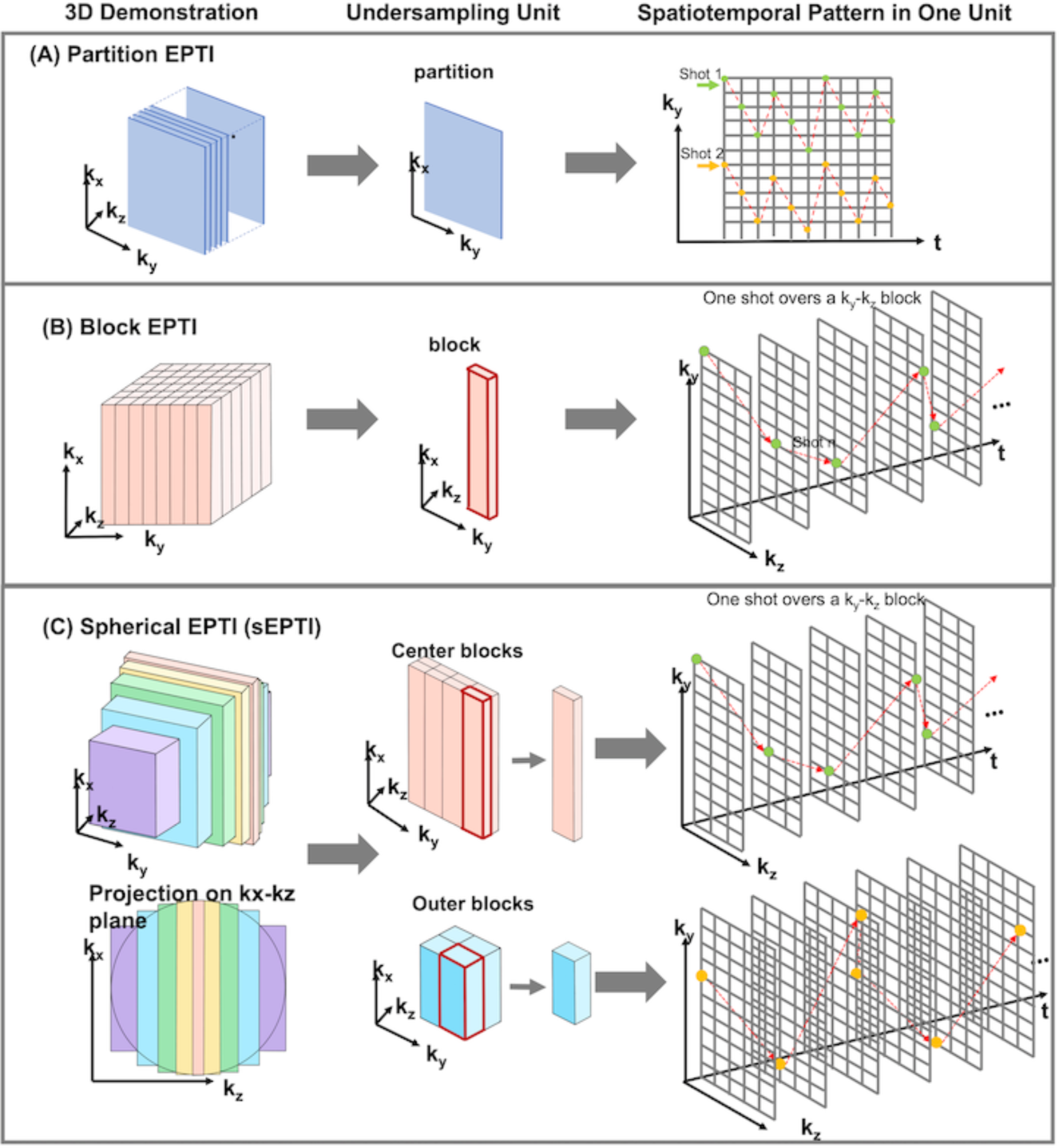
The example sampling pattern for partition EPTI, block EPTI, and spherical EPTI (sEPTI). (A) Partition-EPTI is a direct extension of 2D multi-shot EPTI with undersampling along k_y_ and full-sampling along k_z_. (B) Block-EPTI divides the k_y_-k_z_ plane into blocks at a certain k_y_-k_z_ spacing. Each EPTI-shot samples k_y_ and k_z_ locations within one block using temporal-variant sampling patterns. (C) sEPTI acquires the k-space following a tight spherical envelope. It reduces the number of EPTI-shot through: (1) cutting the k_y_-k_z_ corners; (2) reducing k_x_ extend at outer k_y_-k_z_ locations following a spherical envelope which reduces the echo-spacing and enable a larger k_y_-k_z_ block size. Each EPTI-shot samples k_y_ and k_z_ locations within one block using a pseudo random sampling patterns.

To leverage the undersampling in both phase encoding (k_y_ and k_z_) directions, Dong et al developed a k_y_-k_z_-t undersampling trajectory (Figure 1B), referred to as block-EPTI^31^. This strategy divides the 3D k-space into rectangular blocks at a certain k_y_-k_z_ spacing. Each EPTI-shot samples k_y_ and k_z_ locations within one block using temporal-variant sampling patterns for less-coherent sampling. The acquisition time of block EPTI is the product of the number of shots (same as the number of blocks) and the TR.

#### 2.1.2. Spherical EPTI (sEPTI) trajectory

In this work, Spherical EPTI (sEPTI) was developed to provide further improvement in the acquisition speed as well as robustness to field imperfections (achieved via increased temporal incoherency). Similar to block-EPTI, sEPTI (Figure 1C) also acquires k-space in blocks, but it reduces number of EPTI shots through: (1) cutting the k_y_-k_z_ corners; (2) reducing k_x_ extend at outer k_y_-k_z_ locations following a spherical envelope^34,35^ which reduces the echo-spacing and enable a larger k_y_-k_z_ block size. By employing spherical coverage, a maximum reduction of scan time of 1 − *π*/6 ≈ 0.48 is possible^36^.

Ramp sampling is also utilized to further improve the sampling efficiency by reducing the echo spacing as well as the TR. When compared to no-ramp sampling, the inclusion of maximum ramp sampling reduces the echo spacing by 25% for 1 mm resolution (Supporting Information Figure S1) and almost 30% for 0.75 mm resolution with a maximum gradient (Gmax) 40 mT/m and a maximum slew rate (Smax) 120 mT/m/ms.

### 2.2 Image reconstruction

#### 2.2.1 Low-rank subspace reconstruction

The low-rank subspace reconstruction is used to exploit the spatiotemporal correlation along the TE^31,37^. The spatiotemporal image *I* ∈ ℂ^*N_r_*×*N_t_*^ contains *N*_*r*_ voxels and *N*_*t*_ echoes. The high correlation along TEs enables partial separability of *I* = **B**_p_**UΦ**, where **B**_p_ is the matrix formation of B_0_ inhomogeneity induced phase evolution over TEs, **U** ∈ ℂ^*N_r_*×*L*^ is the spatial coefficients representing the spatial variation, **Φ** ∈ ℂ^*L*×*N*_*t*_^ is the real-valued temporal subspaces containing the temporal dynamics, and *L* is the rank. By separating the phase evolution **B**_p_ from **UΦ**, **UΦ** contains only the background phase, which is lower in rank compared to *I*. The image reconstruction in general can be expressed as

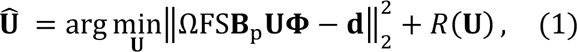

where **d** is the acquired data, Ω is the EPTI undersampling mask, F is the Fourier transform, S is the coil sensitivity, **B**_p_ = *e*^*j*2*πγB*_0_*T*^ is the phase with a static field inhomogeneity map *B*_0_ over a vector of TEs, and *R*(·) is transform domain constraint on the spatial coefficients, which was selected as Wavelet constraint for this work.

#### 2.2.2 Iterative rank-shrinking B_0_ update

Equation 1 intrinsically requires an accurate B_0_ map for high-quality image reconstruction. A common solution is to obtain a low-resolution B_0_ map (*B*_0,low_) by acquiring low-resolution multi-echo calibration data, which can be done in 20 seconds or less. However, directly using *B*_0,low_ for reconstruction can introduce artifacts in areas with spatially-rapidly-changing B_026_. Additionally, the actual B_0_ can change between the calibration scan and the EPTI acquisition due to system drift and motion, which can result in a spatially varying change up to ±20 Hz^38^ in B_0_. To resolve the issue, Dong et al developed a data-driven B_0_ update algorithm^27^, which provides a high-resolution B_0_ map from the EPTI data using *B*_0,low_ as an initial guess. The high-resolution B_0_ is obtained through:

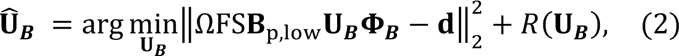

where **d** is the acquired data, **Φ**_***B***_ is temporal subspace containing both T2* decay and phase evolution with off-resonance [−50, 50] Hz, **B**_p,low_ is the phase induced by *B*_0,low_, and **U**_***B***_ is the spatial coefficients to be solved. The residual phase *e*^*j*2*πγB*_0_*T*^ that is not contained by **B**_p,low_ can be obtained from the phase of the reconstructed images **U**_***B***_**Φ**_***B***_, and the B_0_ map for following data reconstruction is *B*_0_ = *B*_0,*low*_ + Δ*B*_0_.

This data-driven update was shown to improve the accuracy of B_0_ and the quality of reconstruction. However, **Φ**_***B***_ containing phase usually doubles the rank of **Φ**, making the B_0_ update an ill-posed problem, and leading to noisy outcomes or increased error for higher-resolution EPTI. To obtain an accurate and high-quality B_0_ map, in this work, an improved iterative rank-shrinking B_0_ updating algorithm was developed (Figure 2A), which updates the B_0_ iteratively in *N* iterations. In iteration *i* ∈ [1, *N*], the B_0_ updating is performed as

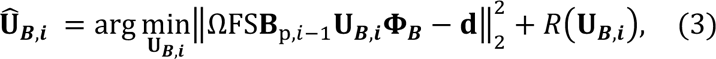

**Figure 2:**
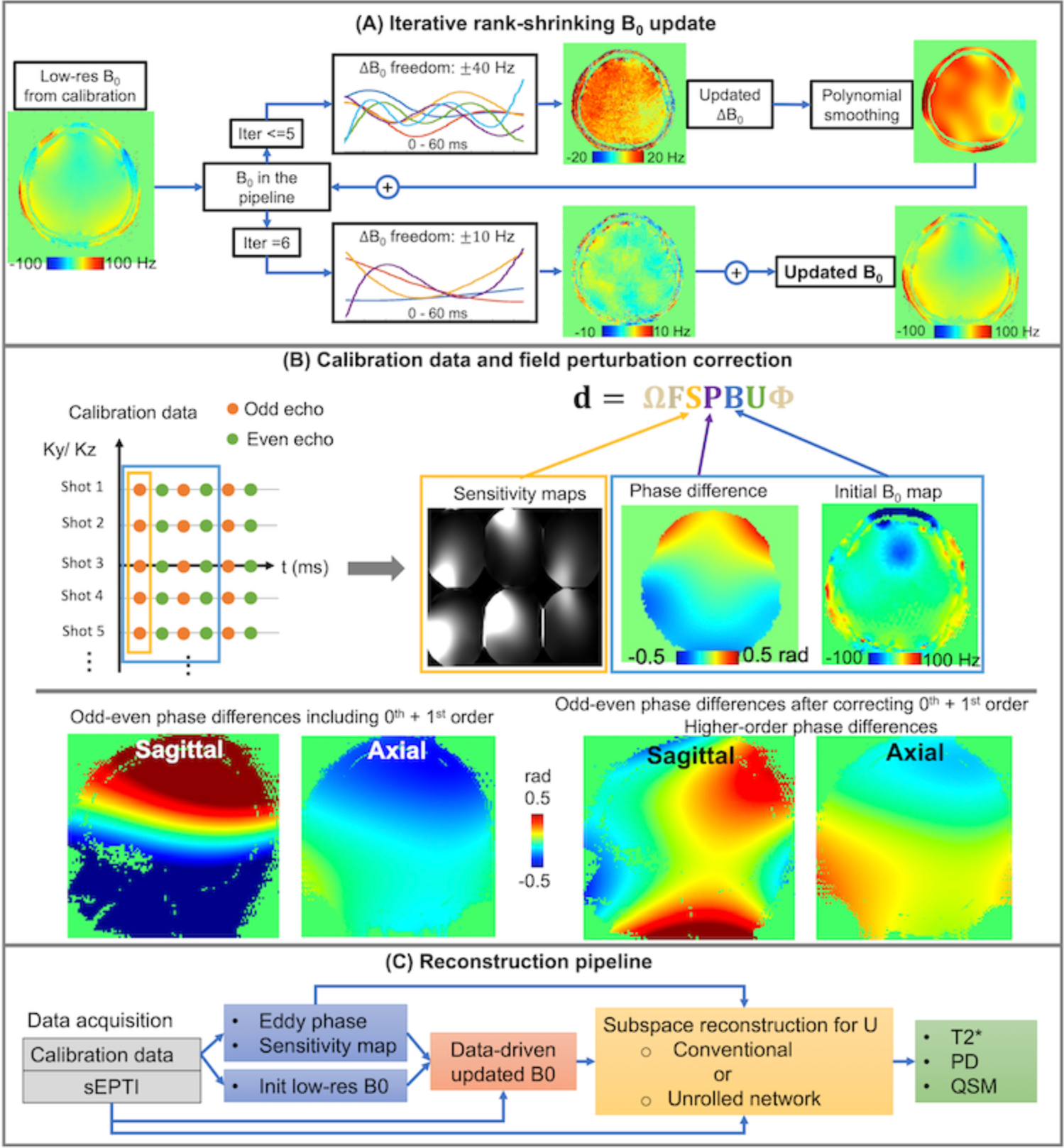
The reconstruction pipeline. (A) the iterative rank-shrinking B_0_ update pipeline to obtain a smooth B_0_ map with sharp edges on the tissue air interface. An initial B_0_ map was obtained from calibration data. For the first 5 iterations, a ΔB_0_ within ±40 Hz was captured and smoothed through the polynomial fitting, and an updated B_0,i_ = B_0,i-1_ + ΔB_0,i_ was achieved in each iteration; in the last iteration, a ΔB_0_ within ±10 Hz was captured with reduced rank to retain the sharp edge of B_0_. (B) the higher-order odd-even phase correction. The correction is achieved by incorporating a phase map representing the phase differences between odd and even echoes into the subspace reconstruction. The phase difference map is obtained from a low-resolution multi-echo calibration data fully sampled in k_y_-k_z_-t dimension. (C) the integrated reconstruction pipeline incorporating all the correction steps.

where **B**_p,*i*−1_ is the phase induced by *B*_0,*i*−1_ estimated from iteration *i* − 1, and **U**_***B***,***i***_ is the spatial coefficients in current iteration. A high-order polynomial fitting is applied to Δ*B*_0,*i*_ for smoother outcome. The updated B_0_ map in iteration *i* is *B*_0,*i*_ = *B*_0,*i*−1_ + Δ*B*_0,*i*_. In the first iteration (*i* = 1), the initial phase **B**_p,0_ is generated from the low-resolution B_0_ map (*B*_0,0_ = *B*_0,low_). In the last few iterations, less off resonance ([−10, 10] Hz) as well as lower rank is used for better conditioning, and polynomial fitting is excluded to preserve the sharp edge of B_0_. The final B_0_ map *B*_0,*N*_ is used to generate the phase **B**_p_ in Equation 1.

#### 2.2.3 Correction of odd-even phase differences

The artifacts from phase differences between odd and even echoes caused by gradient delays and eddy current are a long-standing challenge for EPI-based acquisitions. To correct the gradient-delay-induced 1^st^ order phase difference in readout direction and a 0^th^-order eddy current phase difference between the odd and even lines, 1D readouts at center k_y_-k_z_ are usually acquired in both polarities as odd-even navigators. However, these 1D navigators are not enough to correct the residual higher-order phase differences, which can lead to additional artifacts. To compensate for this issue, we utilized a high-order phase correction pipeline proposed by Dong et al^27^ and generalized it for this 3D EPTI problem. Here, a 3D phase difference map ***p*** ∈ ℂ^*N_r_*×1^ between the odd and even echoes is estimated from the same low-resolution calibration data used for B_0_ initialization and extrapolated by 3^rd^-order polynomial fitting. In the reconstruction model, **P** = [***p***, **0**, ***p***, **0**, ⋯] ∈ ℂ^*N_r_*×*N*_*t*_^ is added to Equation 1, resulting in the following equation:

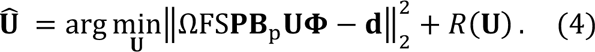

The field correction algorithm is demonstrated in Figure 2B. The illustration of the entire reconstruction pipeline incorporating all the pieces is in Figure 2C.

### 2.3 SNR boosting using physics-informed unrolled network

SNR is always a challenge for T2* and QSM quantification because they both require images at long TE with reduced SNR. Moving to submillimeter resolution using short scan time significantly aggravates this problem. In recent years, deep learning (DL) methods using unrolled network have been widely used in MRI image reconstruction for reduced artifacts and increased SNR^39–41^. Such methods incorporate the DL network into the MRI forward imaging model as a regularization term and solve the inverse problem by alternating between the data consistency (DC) term and the regularization term. Compared to pure data-driven methods, unrolled network leverages MRI physics, and thereby achieves more robust performance and requires less training data^33^.

In this work, a subspace-based unrolled network was incorporated to boost the SNR of sEPTI to achieve high-SNR T2* and QSM quantification at submillimeter resolution in a short single average scan. Instead of Equation 4, the image reconstruction is now performed using the MoDL^42^ architecture as

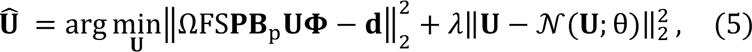

where 𝒩(·) is the convolutional neural network (CNN)-based denoiser with trainable parameters θ. The network was trained in a supervised manner, with the single-average undersampled sEPTI data as input and the reconstructed spatial coefficients **U**_GT_ from multi-average sEPTI data as ground truth (GT). A *l*_1_ loss was used as

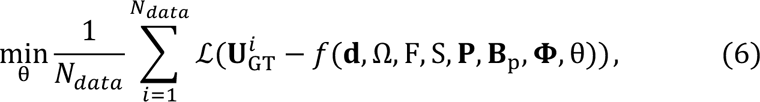

where *N*_*data*_ is the number of training datasets, 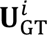 is the *i*-th GT spatial coefficients, *f*(·) denotes the output of the unrolled reconstruction from Equation 5, and ℒ(·) is the loss between the GT and the network output. The image reconstruction pipeline with unrolled network is illustrated in Figure 3.

**Figure 3:**
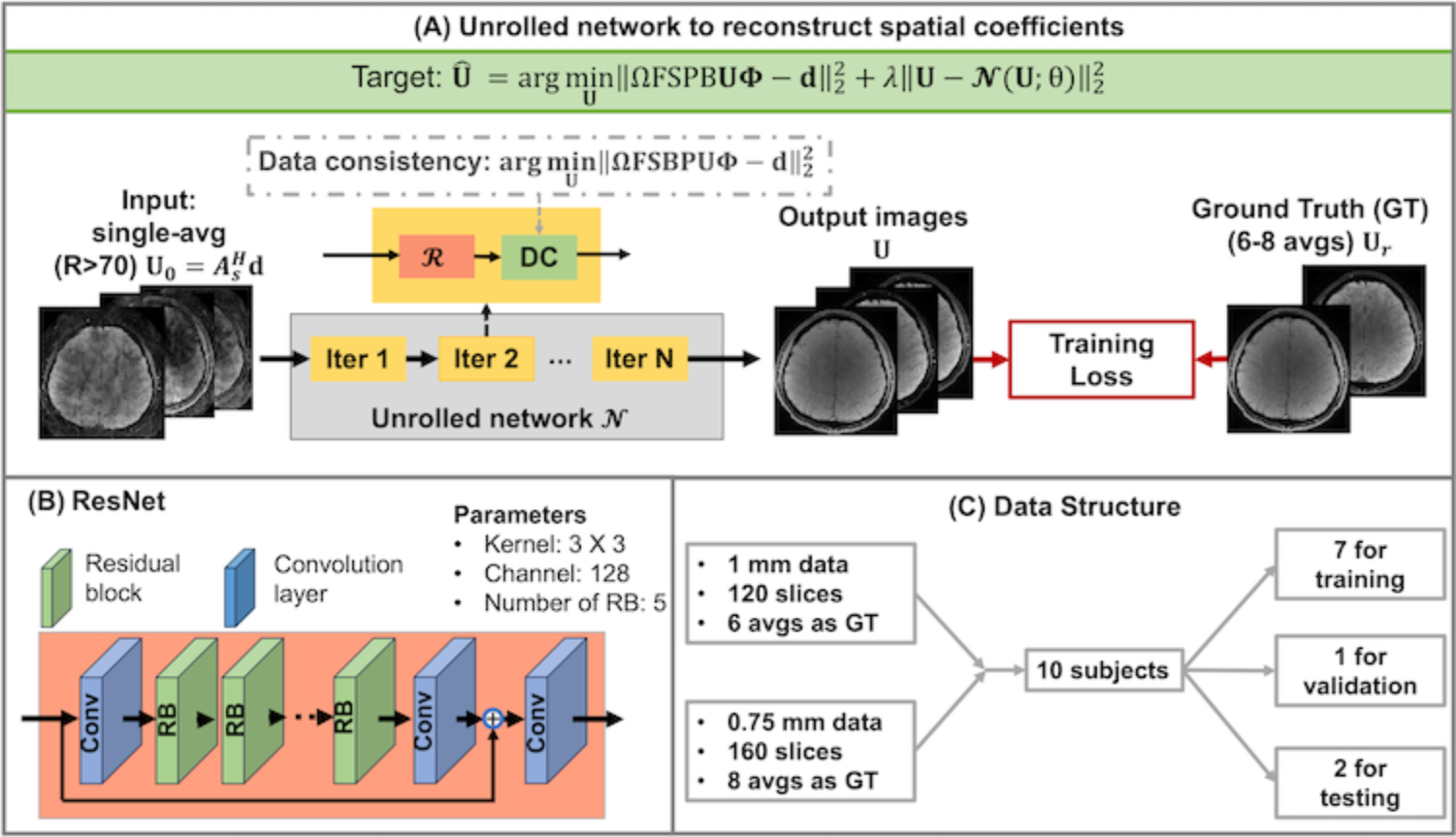
Demonstration of the unrolled network. (A) the structure of the unrolled network. The input is the initial guess from single-average data, and the target is the 6-8 average coefficient maps. The network is a ResNet with hyperparameters specified in (B). (C) Ten subjects were recruited and the data from 7 of them were used for training. Each subject was scanned for 1-mm data with 6 averages and 0.75 mm data with 8 averages. For 1-mm data, each dataset contains 120 slices with good brain structure; for 0.75mm data, each contains 160 slices.

## 3 Methods

### 3.1 Sequence configuration

A 3D gradient echo sequence is used to generate the T2* and QSM contrast. A 4.4-ms SLR pulse was designed for non-selective excitation with a null excitation at the main fat peak, achieving water-only excitation without the need of additional fat-saturation module for a shorter minimum achievable TR. The shape and frequency response of the SLR pulse are shown in Supporting Information Figure S2. Each excitation pulse is followed by one sEPTI-shot.

### 3.2 Imaging protocols

All studies were performed on a 3T system (Premier, GE Healthcare, Milwaukee, USA) with a commercial 48-channel head coil. Images were acquired in Sagittal orientation with following sequences:

1. A fully sampled k_y_-k_z_-t calibration sequence was acquired to estimate the coil sensitivities and the odd-even field perturbation maps, as well as the initial low-resolution B_0_ map. It has FOV = 240×240×216 mm^3^, matrix size = 120×50×40, TR = 10 ms, TEs = 6.0/6.6/7.2/7.8 ms, flip angle = 8°, and scan time = 20 seconds.
2. A reference EPTI protocol at 1 mm was acquired to provide the reference values for T2* and QSM. It fully samples the central k_y_-k_z_ area and undersamples the outer k_y_-k_z_ area with R=2×2. The FOV = 240×240×216 mm^3^, matrix size = 240×240×216, TR = 40 ms, TEs = 5-36 ms, echo spacing = 1.05 ms, flip angle = 16°, and scan time = 10 minutes.
3. 3D partition-EPTI, block-EPTI, and sEPTI were acquired at 1-mm resolution with the following parameters: FOV = 240×240×216mm^3^, matrix size = 240×240×216, TR = 60 ms, TEs = 5-56 ms, flip angle = 20°. For partition-EPTI, the in-plane acquisition used 2D 5-shot EPTI trajectory with an undersampling factor *R* = 48 and scan time = 65 seconds. For block-EPTI, the block size in k_y_-k_z_ plane is 12×4, resulting in *R* = 48 and scan time = 65 seconds. For sEPTI, larger block sizes are used at the outer k_y_-k_z_ positions where the echo-spacing is reduced from shorter k_x_ transversal. Here, block size progressively increases in 4 steps from 12×4 in central k-space region to 12 x 5, 12x 6, and 12x7 at the outermost k-space region, resulting in an averaged *R* ≈ 69 and scan time = 45 seconds. In all cases, the under-sampling factor is calculated based on the time saving in relation to the full k-t sampling scan where acquisition is performed for one k_y_-k_z_ position per TR. The echo spacings for partition EPTI and block EPTI are both 1.05 ms, while for sEPTI, the echo spacing ranges from 1.05 ms of central shots, 0.90 ms, 0.75 ms, and 0.6 ms for the outermost shots (Supporting Information Figure S3), resulting in incoherent echo times through the acquisition.
4. 3D sEPTI was acquired at 0.75-mm resolution with parameters: FOV = 240×240×216mm^3^, matrix size = 320×320×288, TR = 60 ms, TEs = 5-56 ms, flip angle = 20°, the echo spacings are 1.20 ms of the central shots, 1.08 ms, 0.94 ms, 0.78 ms, and 0.62 ms of the outmost shots, and scan time = 90 seconds.

The detailed parameters for all sequences are listed in Supporting Information Table S1.

### 3.3 Simulations

Numerical simulations were performed using a set of 1-mm proton density (PD), T2*, sensitivity, B_0_, and odd-even phase difference maps obtained from the reference EPTI data. In the simulations, the odd-even phase difference maps were used to mimic the field imperfections. The imaging parameters used to simulate partition EPTI, block EPTI, and sEPTI were the same as in Section 3.1. To evaluate the robustness to field perturbations for different trajectories, image reconstruction was performed without and with the odd-even phase difference map incorporated, using Equation 1 and Equation 4, respectively.

### 3.4 Phantom experiments

The phantom study was performed on the ISMRM/NIST MRI system phantom^43,44^ to validate the T2* mapping accuracy. In the phantom study, the calibration data was performed first, followed by the 1-mm reference EPTI, partition EPTI, block EPTI, and sEPTI. The Bland-Altman plots were used to analyze the agreement between the T2* values from reference EPTI and undersampled EPTI methods.

### 3.5 In vivo experiments

The in vivo study was approved by the institutional boards. Written informed consent was obtained from all participating subjects before scanning. Fifteen healthy volunteers (aged 22 to 60 years) without known neurological diseases were recruited for the study. Data from 10 volunteers were dedicated to the training of the unrolled network. The ground truth (GT) high-SNR EPTI data were acquired as multiple averages of sEPTI (6 averages for 1-mm sEPTI, and 8 averages for 0.75-mm sEPTI) rather than a single EPTI scan at a low acceleration factor to avoid artifacts from subject motion and system drift that can be present during long scans. For the other 5 volunteers, the calibration data, reference EPTI (1 mm), partition EPTI (1 mm), block EPTI (1 mm), and sEPTI (1 mm and 0.75 mm) were acquired using the same parameters as in Section 3.1.

To test the generalizability of the proposed sEPTI technique, two pediatric patients with epilepsy, whose T2* and QSM values are different from the healthy adult population, were recruited and the 1-mm sEPTI were acquired.

### 3.6 Network and training details of unrolled reconstruction

The neural network in Equation 5 was implemented using a ResNet^45^ structure (Figure 3). The ResNet consisted of a layer of input and output convolution layers and 5 residual blocks. Each residual block consists of two convolutional layers, in which the first layer is followed by a rectified linear unit, and the second layer is followed by a constant multiplication layer. All layers have a kernel size of 3 × 3 and 128 channels. For each batch of data, the network was unrolled for 10 iterations.

During the acquisition of GT, it is possible to have bulk motion and background phase drifting through multiple averages. To ensure that our generated GT is of good quality, each average was reconstructed using Equation 4 separately. Subsequently, the spatial coefficients for each average were co-registered to the first average using FSL, and a low-resolution background phase difference^46^ of the spatial coefficients was also corrected to match that of the first average. Finally, a complex average^47^ was performed on the spatial coefficients of all averages as GT.

The unrolled reconstruction is performed in a 2D manner. For the acquired sEPTI data, a 1D NUFFT (non-unform sampling because of the ramp) was first performed in k_x_ direction to transform the data into a hybrid x-k_y_-k_z_-t format. The reconstruction was then performed for each x-slice. Data from 7 out of 10 subjects were used for training, 1 for validating, and 2 for testing. For each subject, 1-mm sEPTI data provides 120 slices containing main brain structures, while 0.75-mm data provides 160 slices, resulting in a total (120 + 160) × 7 = 1960 slices for training. The network was trained on 1-mm data only, 0.75-mm data only, and both data, respectively, and the performance of different networks were evaluated on the validating and testing datasets through structural similarity index measure (SSIM) and root mean square error (RMSE) against the GT.

## 4 Results

### 4.1 Simulations

The simulation results are demonstrated in Figure 4. When incorporating the odd-even echo phase differences in both the generation and reconstruction of the images, partition EPTI, block EPTI and sEPTI all showed good image quality with low RMSE. Partition EPTI (R = 48) produces the highest errors and the most noise since it concentrates the undersampling only in the k_y_ direction, resulting in an elevated g-factor. Block EPTI (R = 48) provides the lowest error with mean relative RMSE = 4.8%. For sEPTI (R = 68), a slightly higher error is observed when compared to block EPTI, with mean relative RMSE = 5.6%, due to the cut-off of k-space corners and the 1.4× higher acceleration in acquisition (45 s vs 65 s).

**Figure 4.**
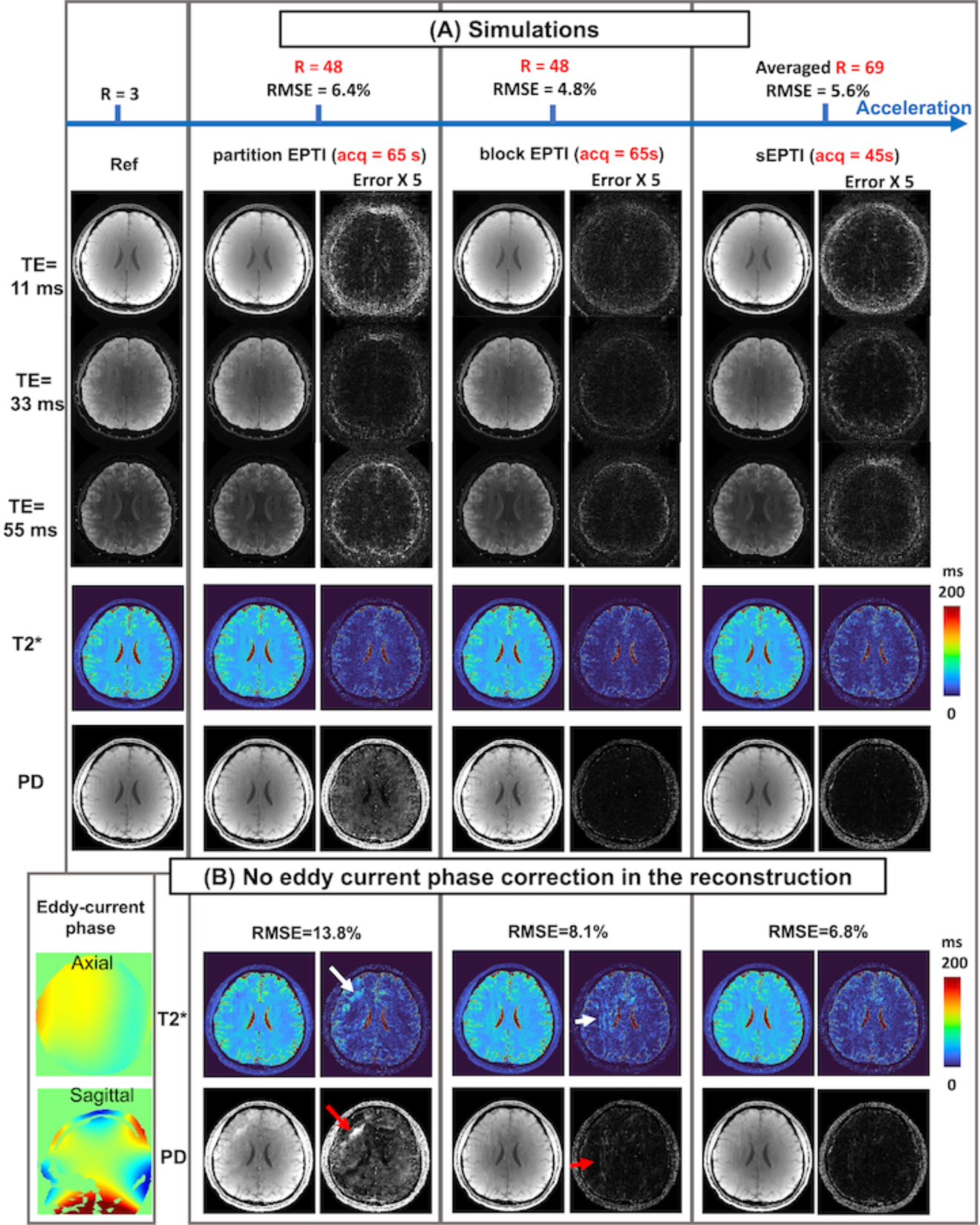
Simulation results. (A) The eddy current phase differences between odd-even echoes were incorporated in both generation and reconstruction. Partition EPTI (R = 48) showed higher noise. Block-EPTI achieved improved image and mapping qualities. sEPTI provided similar image and mapping quality in 70% of the acquisition time. (B) The eddy current phase differences were incorporated in the generation but were excluded from reconstruction to mimic the realistic cases. The eddy current phase is shown in axial and sagittal views. Without correcting the eddy-current phase in the reconstruction, partition EPTI and block EPTI showed decreased image quality (labeled by white and red arrows), while sEPTI is the most robust with similar RMSE as in (A). RMSE: root mean square error

Figure 4B examines the reconstruction errors with the absence of higher-order odd-even phase differences in the reconstruction. Here, partition EPTI shows more than 2x higher error (mean relative RMSE = 13.8%) with strong aliasing artifacts when compared to the results in Fig4A. Similarly, Block EPTI shows increased errors as well as aliasing artifacts (mean relative RMSE from 4.3% to 8.1%). On the other hand, sEPTI with its increased spatiotemporal incoherency, is the most robust to the odd-even phase differences and potential residual phase errors in data, producing the best image quality and a mean relative RMSE from 5.6% to 6.8%.

### 4.2 Phantom results

Imaging results from the T2 layer within the NIST phantom were used for analysis. There are 14 spheres with T2* values varying from 10-600 ms as measured by the reference EPTI sequence. Eleven spheres with T2*<200 ms were included in the analysis, as displayed in Figure 5. Partition EPTI shows noisy and aliased PD and T2* maps and large differences in T2* values compared to the reference with a 19.1-ms standard deviation (SD). Block EPTI provides improved PD and T2* maps. The SD of the T2* difference between block and reference EPTI is 6.5 ms. sEPTI, with a shorter scan time, produces the best PD and T2* maps as well as the most accurate T2* values against the reference with an SD of 3.2 ms.

**Figure 5:**
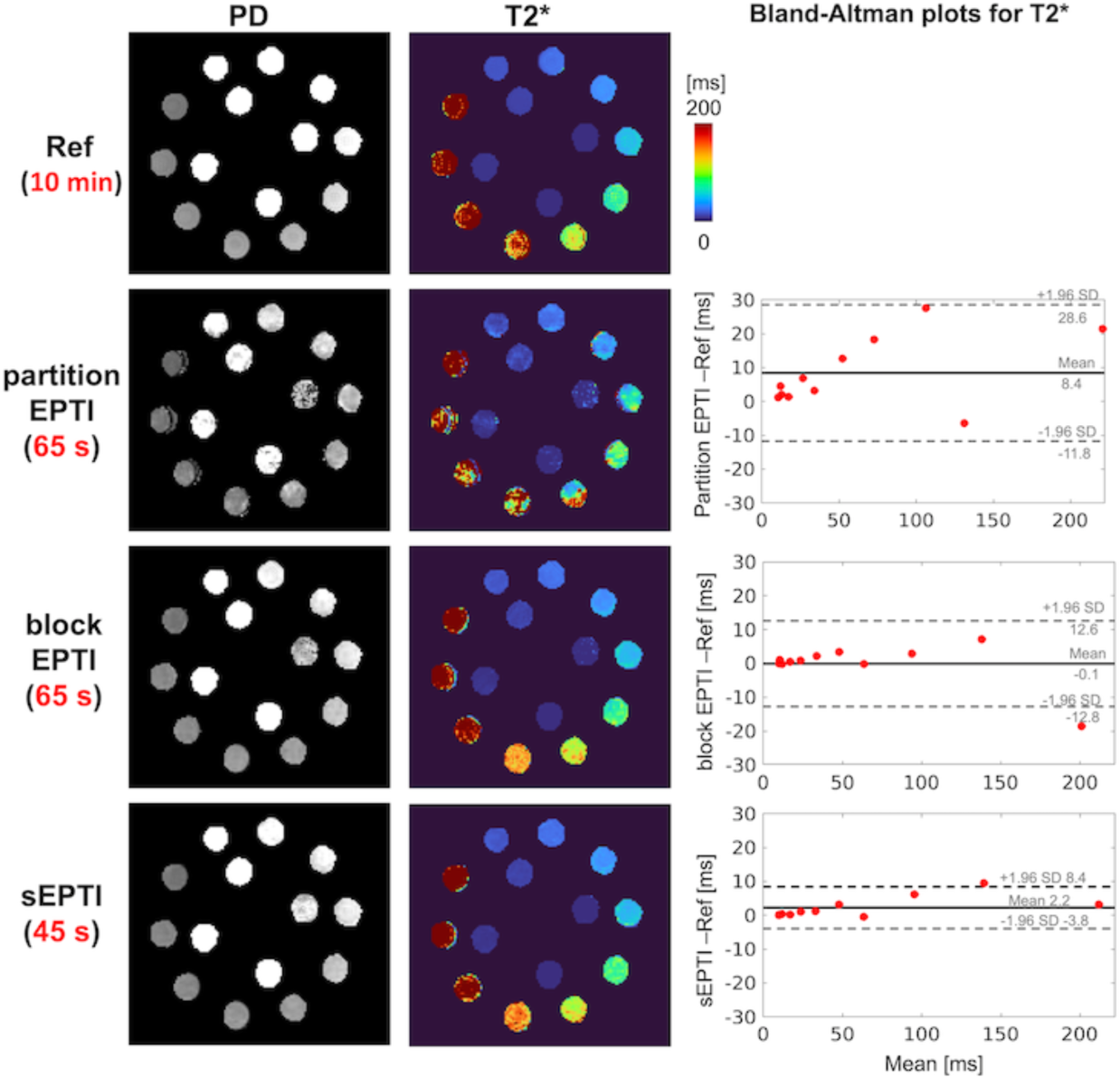
Phantom results. The PD and T2* maps from partition EPTI are noisy and subject to artifacts. For block EPTI, the results are largely improved. sEPTI produces the best PD and T2* maps. The Bland-Altman plots show the T2* values from sEPTI have the smallest standard deviation (SD) compared to the reference.

### 4.3 In vivo results

#### 4.3.1 The iterative rank-shrinking B_0_ update and higher-order odd-even phase correction

Figure 6 illustrates the improvement in the image quality with higher-order odd-even phase correction and iterative rank-shrinking B_0_ update. Figure 6A shows the estimated B_0_ map, T2* weighted image at TE = 33 ms, and PD and T2* maps of the 0.75-mm sEPTI data obtained using previous B_0_ update^27^ and with only 0^th^ and 1^st^ order odd-even phase correction. The outcomes are noisy due to the high resolution and high undersampling rate with visible artifacts from eddy currents. By incorporating the higher-order odd-even phase into the reconstruction pipeline, the eddy-current-related artifacts are extensively reduced, but the outcomes are still noisy (Figure 6B). If directly smoothing the B_0_ using polynomial approximation, the outcome possesses less noise but more aliasing artifacts because the B_0_ map is oversmoothed and inaccurate, particularly at the edge of the tissue-air interface (Figure 6C). The proposed iterative rank-shrinking B_0_ update pipeline maintains the smoothness within the brain as well as the sharpness at the interface, producing high-quality images without aliasing (Figure 6D).

**Figure 6:**
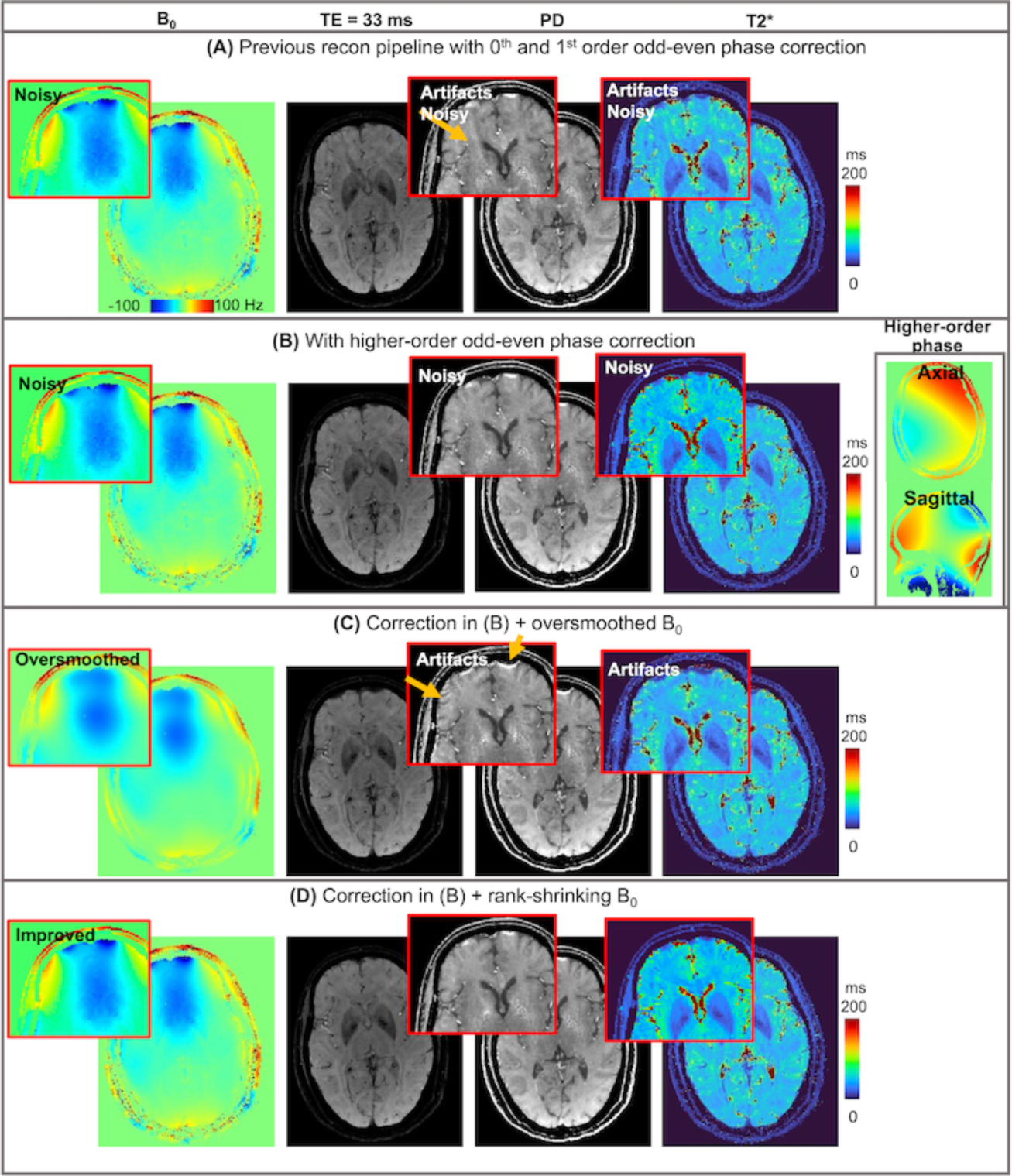
The effect of the higher-order odd-even phase correction and iterative rank-shrinking B_0_ update on a 0.75-mm sEPTI dataset. (A) The B_0_ map, T2*-weighted image, PD, and T2* maps using the previous reconstruction pipeline without the improvements proposed in this work. The images are noisy with strong eddy-current-induced artifacts (yellow arrows). (B) The higher-order odd-even phase correction method can effectively reduce the eddy-current-related artifacts for data, but the outcomes are still noisy due to the noisy B_0_ map. (C) when performing a polynomial fitting to B_0_, the outcomes are less noisy but have artifacts due to inaccurate (oversmoothed) B_0_. (D) By integrating the odd-even phase correction and iterative rank-shrinking B_0_ updating algorithm, the outcome images show high SNR and reduced artifacts.

#### 4.3.2 Image reconstruction with unrolled network

Figure 7 displays the unrolled reconstruction results for test data at 1mm and 0.75mm resolutions, obtained using least-square (LS) reconstruction, wavelet-constrained reconstruction, and unrolled network reconstructions that utilize different training datasets: (1) 1-mm training datasets only (840 slices in total); (2) 0.75-mm training datasets only (1120 slices in total); (3) joint training datasets (1920 slices). For the 1-mm case, the results of all networks showed reduced noise compared to conventional reconstruction using the Wavelet constraint. The network from 1-mm training datasets produced good SSIM (0.962) and low relative RMSE (5.3%) compared to the GT; the network from (unmatched) 0.75-mm training datasets produced lower SSIM (0.958) and higher relative RMSE (5.8%) with visible blurring on the structures (highlighted via white arrows); the network training on joint datasets produced the best SSIM (0.965) and the second to best relative RMSE (5.4%).

**Figure 7.**
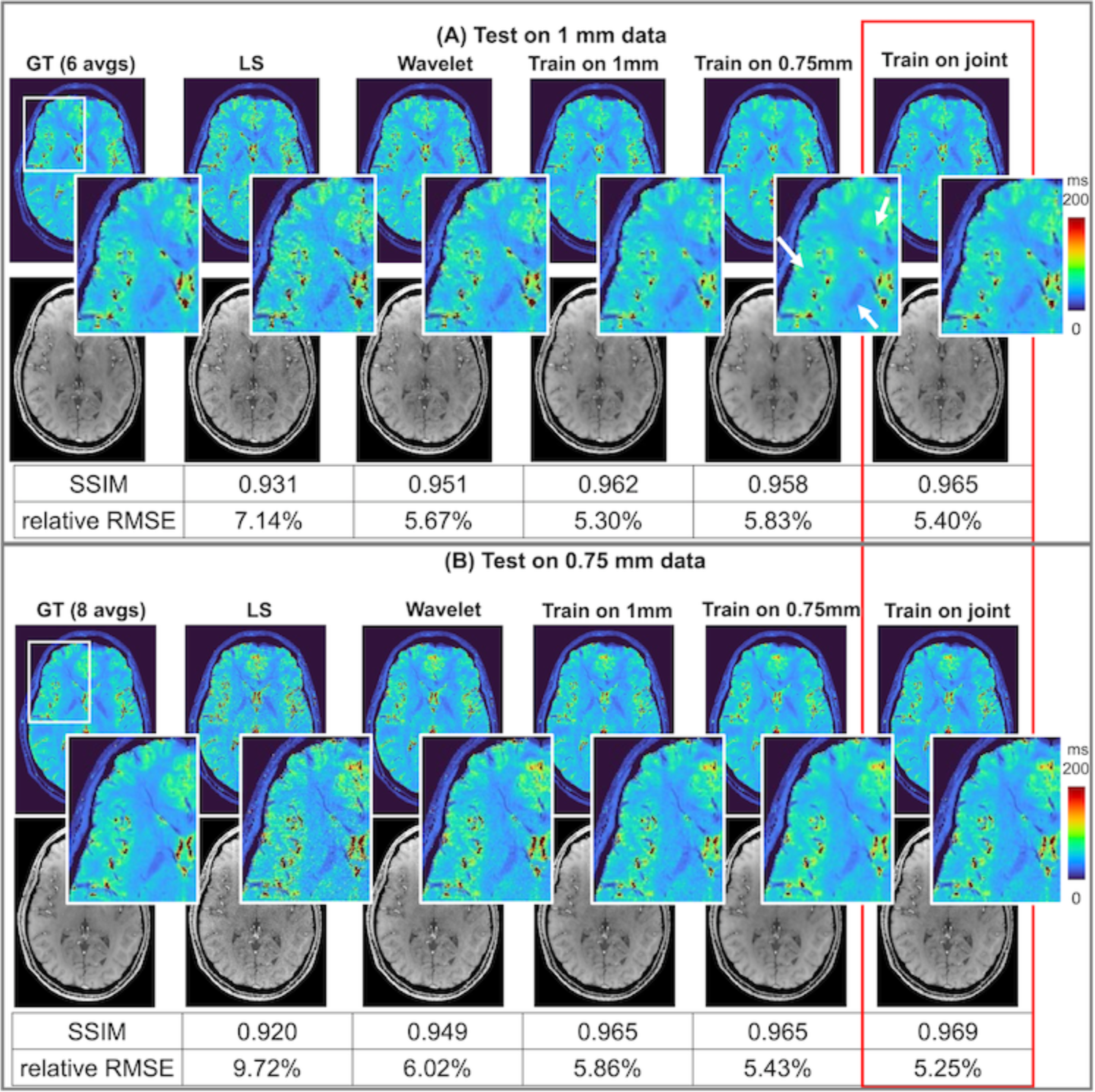
The performance of the unrolled network on the 1-mm test data and 0.75-mm test data. On datasets from both resolutions, the outcomes from unrolled network have significantly improved SNR on the T2* weighted images as well as the T2* maps. The network trained on 0.75-mm data only provides sightly blurry images when applying it to 1-mm test data. The network trained on data of both resolutions provide the highest SSIM for both the 1-mm and 0.75-mm test data.

For the testing on 0.75-mm data, when using conventional reconstruction (LS and Wavelet), the results present increased noise compared to 1-mm test data due to the increased resolution. With unrolled networks, the noise level was significantly reduced. The network training on joint datasets produced the best SSIM (0.969) and the lowest relative RMSE (5.3%).

#### 4.3.3 In vivo experiments

Incorporating all the advances in sampling design and reconstruction, sEPTI produced high-quality whole-brain PD, T2*, and QSM maps at 1 mm resolution within 45 seconds (Figure 8). As a comparison, block EPTI, with a 65-second acquisition time and standard B_0_ update as performed in Dong et al^31^, shows smearing artifacts on PD, T2*, and QSM maps. The 45-second sEPTI produced cleaner images in higher SNR with 1.4× acceleration.

**Figure 8.**
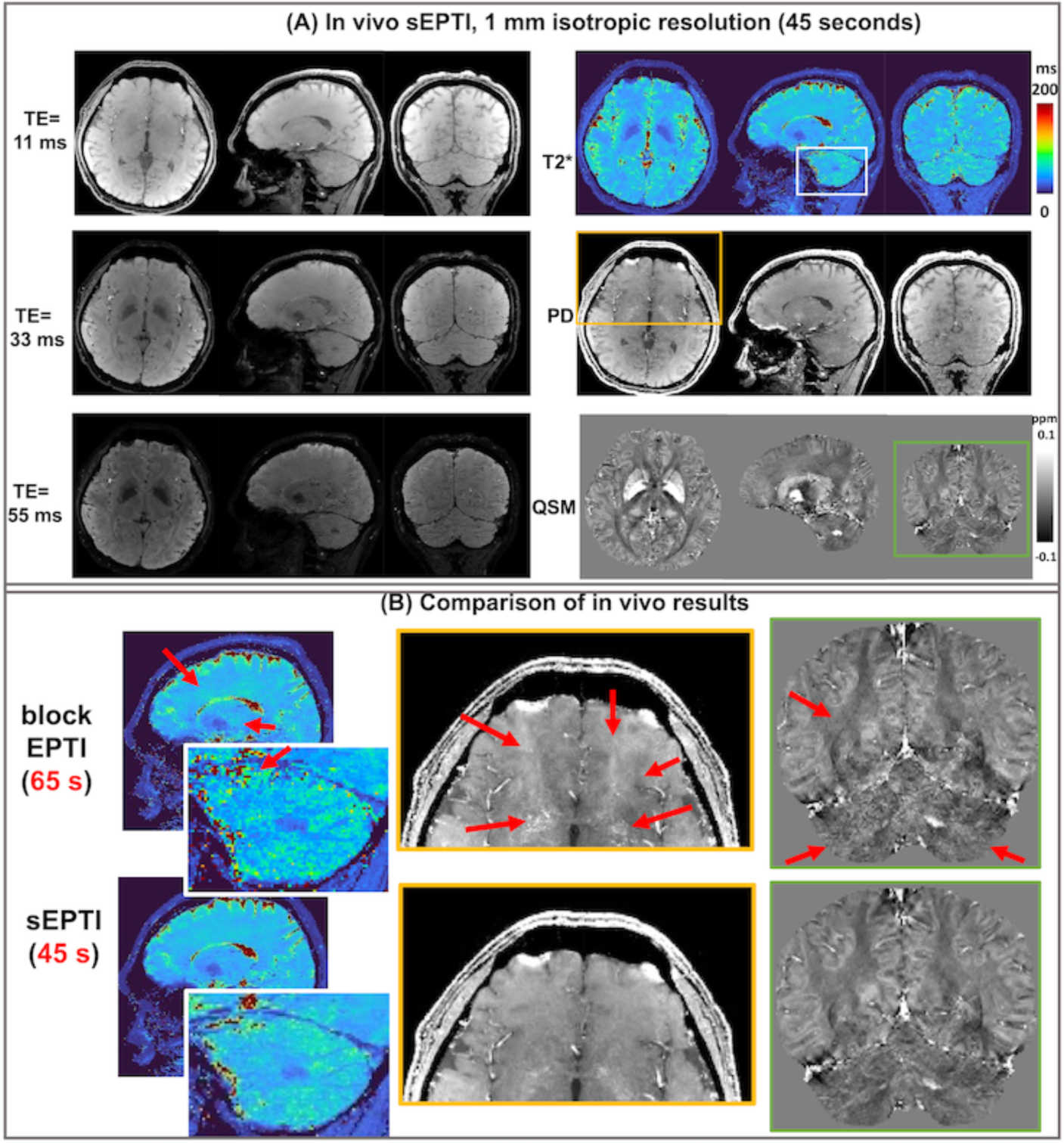
In vivo demonstration. The T2*-weighted images at 11, 33, 55 ms and PD, T2*, and QSM maps were shown for sEPTI in axial, coronal, and sagittal views. The comparison of different EPTI trajectories was shown for zoomed-in areas. Block EPTI with the previous reconstruction pipeline (without the improvements in this work) tends to have more blurring, smearing, and alaising artifacts in PD, T2*, and QSM, while sEPTI with updated reconstruction pipeline provides clean PD, T2*, and QSM maps with clear structure delineation.

The 0.75-mm sEPTI images acquired in 90 seconds are shown in Figure 9. By leveraging the unrolled network for reconstruction, single-average 0.75-mm images showed comparable SNR to the 1-mm images, but with improved capability in delineating fine-scale structures, as illustrated in Figure 9B with red solid arrows.

**Figure 9.**
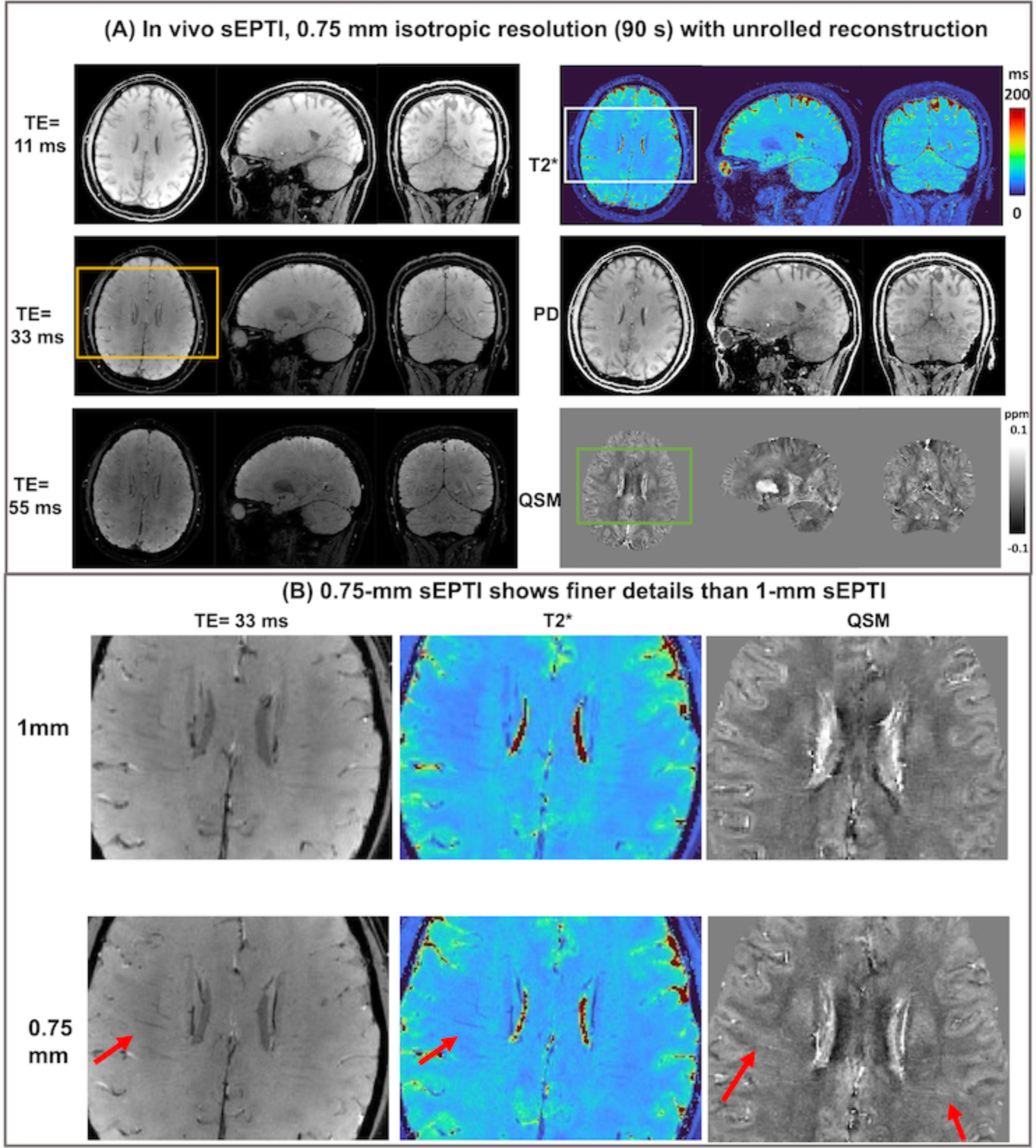
Demonstration of sEPTI with whole-brain coverage at 0.75-mm isotropic resolution. (A) The T2*-weighted images at 11, 33, 55 ms and PD, T2*, and QSM maps were shown in axial, coronal, and sagittal views. (B) a comparison of the 1-mm and 0.75-mm sEPTI on the same subject. On the T2*-weighted images, T2* map, and QSM map, 0.75-mm showed finer structure delineation compared to 1-mm sEPTI.

The generalizability of the developed acquisition and reconstruction pipeline with unrolled network was demonstrated on two pediatric patients with epilepsy. Figure 10 shows the 1-mm sEPTI images of a 9-year-old pediatric patient, whose brain has different T2* and QSM values compared to a healthy adult. In this case, the reconstruction using unrolled network was able to visibly improve the SNR T2* maps compared to the reconstruction using Wavelet constraints, indicating that the acquisition and reconstruction techniques developed in this work can be potentially used for a large variety of patient cohorts.

**Figure 10.**
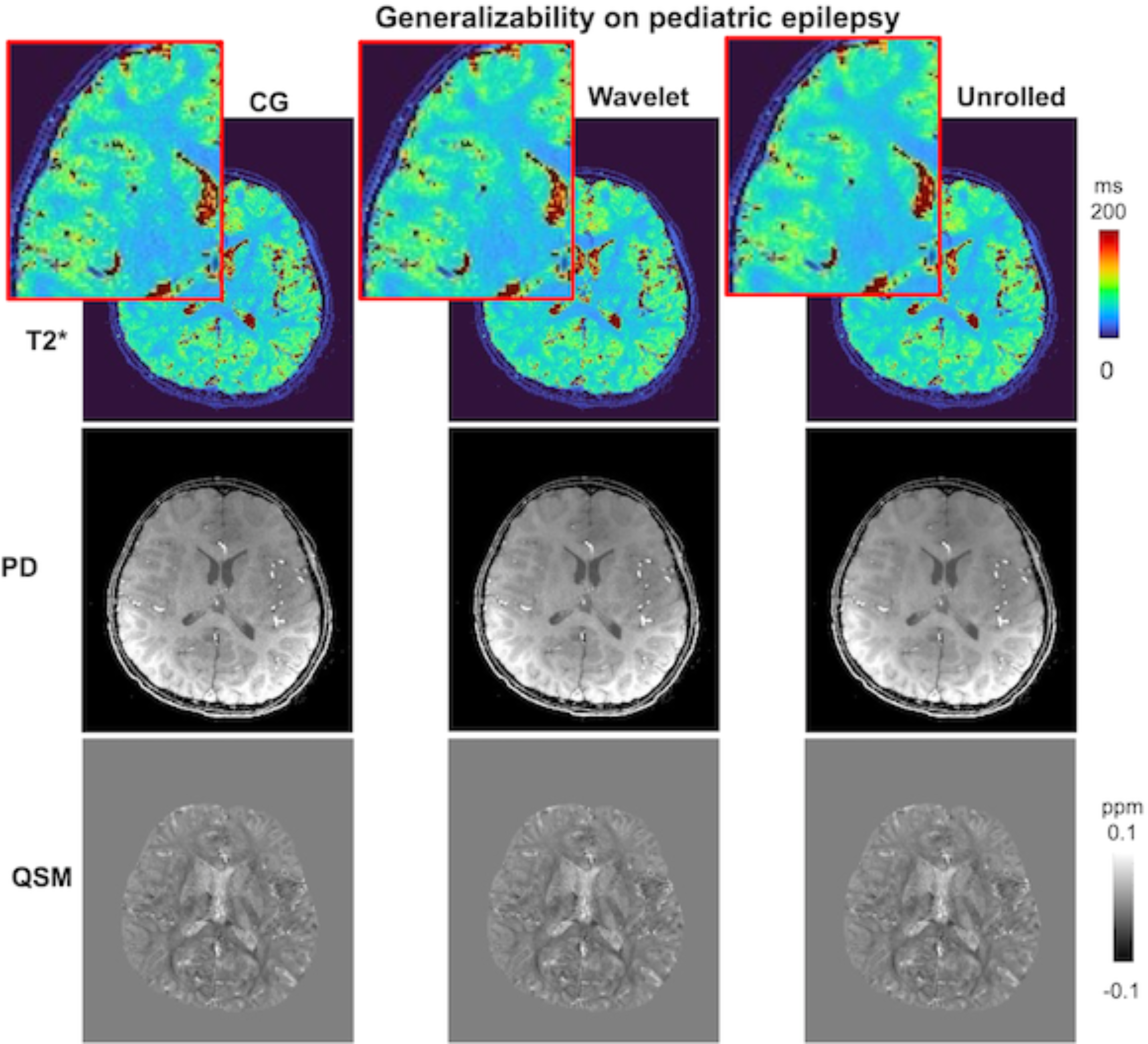
The developed sEPTI and reconstruction pipeline with unrolled network provide high-quality T2* map and QSM for a pediatric subject with epilepsy, where the T2* and QSM values are different compared to the training datasets (healthy adult subjects), indicating the generalizability of the developed pipeline.

## 5 Discussion

In this work, an efficient sEPTI acquisition technique was developed, achieving ≥ 1.4-time acceleration compared to the already-fast EPTI techniques while improving robustness to field imperfections. A possible pitfall is that reducing the cubical k-space coverage to a tight spherical coverage can result in resolution loss due to the cut-off of k-space corner. This was demonstrated in simulations: under ideal conditions, sEPTI produced a slightly higher error than block EPTI, where the minor resolution loss induced only a minor increase in relative RMSE (1.3%). Future evaluation with the recently proposed checkboard test will help the characterization of resolution^48^.

On the other hand, spherical sampling comes with an advantage beyond the increased acceleration. In particular, the varied echo spacing at different k-space locations increases the spatiotemporal incoherency and thus enhances the robustness of sEPTI to field-imperfection-induced artifacts, such as the odd-even phase differences. In the simulations, compared to partition EPTI and block EPTI, sEPTI produced the lowest error when phase errors were not incorporated into reconstruction. In the actual experiments, maximum ramp sampling was utilized to reduce the echo spacing, making the odd-even phase differences a bigger challenge. Under this condition, sEPTI continues to provide cleaner images compared to block-EPTI when the higher-order odd-even phase correction was incorporated for both samples, probably due to sEPTI’s enhanced robustness to the residual field imperfection.

The quality and accuracy of B_0_ maps have an essential influence on the sEPTI image quality. To obtain a high-resolution B_0_ map matching the data, Dong et al^31^. developed a data-driven B_0_ updated algorithm^27^. However, with a higher undersampling rate and higher resolution of sEPTI, the B_0_ update becomes more ill-posed and leads to noisier B_0_ and subsequent noisier PD, T2*, and QSM maps. Performing polynomial fitting on the updated Δ*B*_0_ provides a smoother B_0_, but, at the same time, will oversmooth it in areas with rapid B_0_ spatial variations, resulting in aliased reconstructed images. In this work, based on the B_0_ update approach^31^, we further developed the rank-shrinking B_0_ update algorithm that progressively updates the B_0_ map from the initial guess for an improved outcome. In early iterations, a larger Δ*B*_0_ range and a larger rank are used. The the problem is ill-conditioned and the outcome is noisier, but the polynomial smoothing can improve the results. In later iterations, a smaller Δ*B*_0_ range as well as a smaller rank make the problem less ill-posed, where the polynomial smoothing is no longer needed and thus the sharpness at areas with rapid spatial variation can be kept. The in vivo results demonstrated that the iterative rank-shrinking B_0_ update algorithm significantly improved the SNR and reduced the aliasing of sEPTI. Compared to existing iterative B_0_ and image reconstruction work^49–51^, the proposed approach is well adapted to solve the B_0_ estimation in EPTI, which is a highly ill-posed problem with a high acceleration rate.

The SNR of 0.75-mm sEPTI is 0.75^3^ × e90/45 = 0.6 of that of the 1-mm sEPTI. As illustrated in Figure 7B, the T2* map of 0.75-mm data using Wavelet constraint still contains visible noise in the gray and white matter areas compared to the GT (with 8 average) after utilizing all the advances in the conventional reconstruction developed in this work. After utilizing the unrolled network, the single-average sEPTI data produce high-quality images with improved SSIM and reduced error on test cases. When trained on 0.75-mm data only and applied to 1-mm test data, the outcome from unrolled network tends to be oversmoothed, probably because the 0.75-mm data requires a larger amount of regularization than the 1-mm data. However, when trained on joint datasets, the network outcomes for both 1-mm test data and 0.75-mm test data are of high SNR and sharpness. Therefore, larger sizes of training data with varied features will improve the performance and generalizability of the network.

The network showed promising generalizability. Although trained on datasets containing only healthy adult subjects, the unrolled network was able to produce high-quality, SNR-boosted images on pediatric subjects with epilepsy, whose T2* and QSM values are different compared to healthy adults. This is because the unrolled network mainly performs the regularization of the image while leveraging the MRI imaging physics, and generally has better generalization performance compared to pure data-driven image-domain DL methods^52^.

The developed techniques in acquisition and reconstruction can be flexibly translated to and benefit numerous other techniques and applications. The higher-order odd-even phase correction can help reduce the artifacts in all EPI-based acquisitions^53^. The sEPTI sampling can be easily combined with different sequence structures for multiparametric mapping, diffusion, and fMRI applications.

The technique still contains a number of limitations. First, the sEPTI, although with a short scan time for submillimeter resolution, can still be subject to bulk motion, particularly in motion-prone populations such as the pediatric and the geriatric. Second, the shot-to-shot (TR-to-TR) B_0_ variations, from, e.g., respiratory and bulk motions, are not incorporated into the current reconstruction pipeline. A recently published compact navigator-based motion correction method by our group can estimate the motion and B_0_ variation per TR, which can be a potential solution^54^. In the next step, sEPTI will be applied to patients whose diagnosis can benefit from ultrafast submillimeter T2* and QSM quantifications, such as Multiple Sclerosis patients, to demonstrate its clinical capability further.

## 6 Conclusions

In this work, we developed an sEPTI acquisition technique and a comprehensive reconstruction pipeline with improved B_0_ estimation, higher-order odd-even phase correction, and unrolled network for high-quality submillimeter neuroimaging. The sEPTI showed superior robustness to field imperfections and achieved whole-brain, distortion-free, and blurring-free multi-echo T2*-weighted imaging and T2* and QSM quantification for 1 mm within 45 seconds and 0.75 mm within 90 seconds, promising for future clinical applications.

