## Supporting Information Figure S1 for "Spherical Echo-Planar Time-resolved Imaging (sEPTI) for rapid 3D quantitative T2* and Susceptibility imaging"

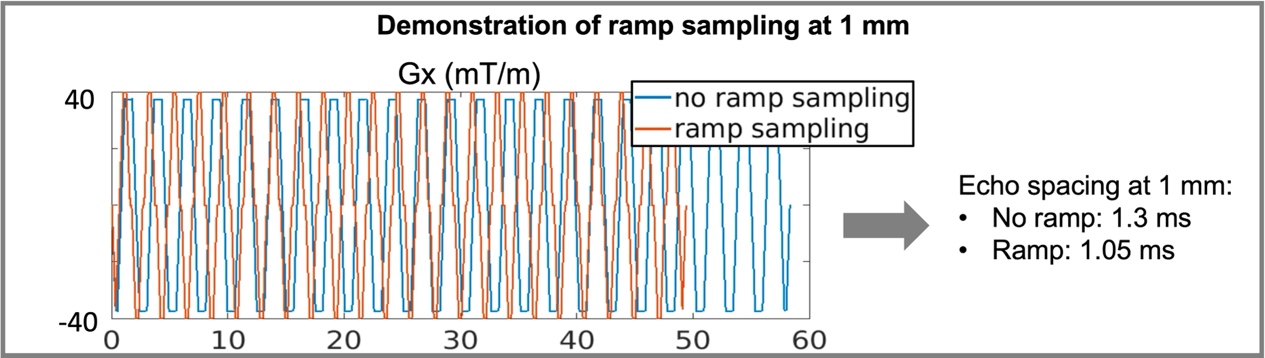

Supporting Information Figure S1: Maximum ramp sampling is used to reduce the echo spacing as well as the total scan time

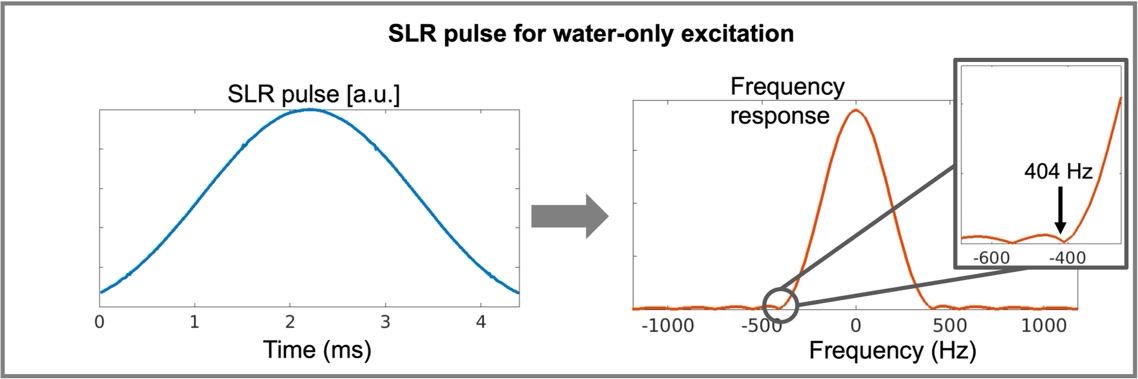

Supporting Information Figure S2: The SLR pulse is designed to suppress fat signal by avoiding main fat peak (~420 Hz on 3T system)

|  | **Calibration** | **Reference EPTI** | **Partition EPTI** | **Block**  **EPTI** | **sEPTI**  **(1 mm)** | **sEPTI**  **(0.75 mm)** |
| --- | --- | --- | --- | --- | --- | --- |
| **FOV (mm^2^)** | 240×240  ×216 | 240×240  ×216 | 240×240  ×216 | 240×240  ×216 | 240×240  ×216 | 240×240  ×216 |
| **Matrix** | 120×50×40 | 240×240  ×216 | 240×240  ×216 | 240×240  ×216 | 240×240  ×216 | 320×320  ×288 |
| **TR (ms)** | 10 | 40 | 60 | 60 | 60 | 60 |
| **TE (ms)** | 6.0/6.6/7.2/7.8 | 5-35 | 5-56 | 5-56 | 5-56 | 5-56 |
| **Echo Spacing (ms)** | 0.60 | 1.00 | 1.05 | 1.05 | 1.05/0.90/  0.75/0.60 | 1.20/1.08/0.94/0.78/0.62 |
| **Flip angle (◦)** | 8 | 16 | 20 | 20 | 20 | 20 |
| **Scan Duration** | 20 sec | 10 min | 65 sec | 65 sec | 45 sec | 90 sec |

Supporting Information Table S1: Imaging protocols

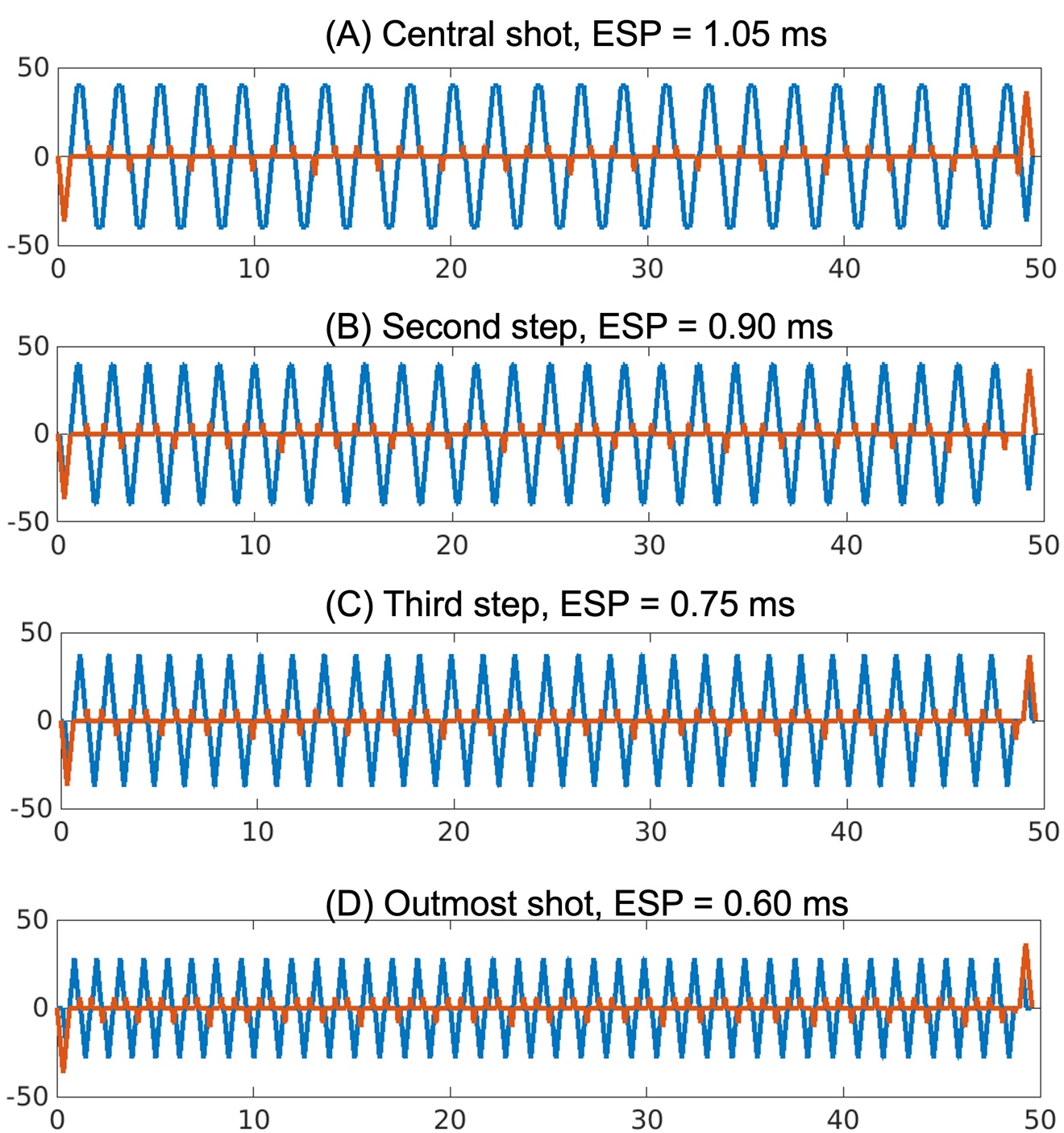

Supporting Information Figure S3: Example gradient waveforms for 1-mm sEPTI design. The echo spacing and block size changes from the center k-space to the outermost k-space in 4 steps.
